# A Network-based Deep Learning Framework Catalyzes GWAS and Multi-Omics Findings to Biology and Drug Repurposing for Alzheimer’s Disease

**DOI:** 10.1101/2021.10.20.465087

**Authors:** Jielin Xu, Yuan Hou, Yadi Zhou, Ming Hu, Feixiong Cheng

## Abstract

Human genome sequencing studies have identified numerous loci associated with complex diseases, including Alzheimer’s disease (AD). Translating human genetic findings (i.e., genome-wide association studies [GWAS]) to pathobiology and therapeutic discovery, however, remains a major challenge. To address this critical problem, we present a <u>**net**</u>work <u>**t**</u>opology-based deep learning framework to identify disease-<u>**a**</u>ssociated <u>**g**</u>enes (NETTAG). NETTAG is capable of integrating multi-genomics data along with the protein-protein interactome to infer putative risk genes and drug targets impacted by GWAS loci. Specifically, we leverage non-coding GWAS loci effects on expression quantitative trait loci (eQTLs), histone-QTLs, and transcription factor binding-QTLs, enhancers and CpG islands, promoter regions, open chromatin, and promoter flanking regions. The key premises of NETTAG are that the disease risk genes exhibit distinct functional characteristics compared to non-risk genes and therefore can be distinguished by their aggregated genomic features under the human protein interactome. Applying NETTAG to the latest AD GWAS data, we identified 156 putative AD-risk genes (i.e., *APOE*, *BIN1*, *GSK3B*, *MARK4*, and *PICALM*). We showed that predicted risk genes are: 1) significantly enriched in AD-related pathobiological pathways, 2) more likely to be differentially expressed regarding transcriptome and proteome of AD brains, and 3) enriched in druggable targets with approved medicines (i.e., choline and ibudilast). In summary, our findings suggest that understanding of human pathobiology and therapeutic development could benefit from a network-based deep learning methodology that utilizes GWAS findings under the multimodal genomic analyses.

## Introduction

Alzheimer’s disease (AD), first described in 1907 by Alois Alzheimer, is the most common type of dementia with gradual cognitive decline and memory loss [1]. AD and AD-related dementias (AD/ADRD) are a major global health challenge and are expected to double in incidence by 2050 [2], affecting 90 million people worldwide [3]. The incidence of AD at the U.S. is expected to double by 2050 [4,5], while the attrition rate for AD clinical trials (2002-2012) is estimated at 99.6% [6]. High-throughput DNA/RNA sequencing technologies have rapidly led to a robust body of genetic and genomic data in multiple national AD genome projects, including the Alzheimer’s Disease Sequencing Project (ADSP) [7] and the Alzheimer’s Disease Neuroimaging Initiative (ADNI) [8]. Genome-wide association studies (GWAS) have identified over 40 AD susceptibility loci [9–12]. Despite this progress in understanding of AD genetic risk, the predisposition to AD involves a complex, polygenic, and pleiotropic genetic architecture; furthermore, massive genetic and genomic data are not effectively explored for AD drug discovery and development yet.

Recent advances in genetics and systems biology have showed that AD is governed by network-associated molecular determinants (termed disease module) of common endotypes or endophenotypes [13,14]. Approaching AD with a simplistic single-target approach has been demonstrated effective for developing symptomatic therapies but ineffective when attempted for disease modification [13]. Therapeutic approaches by specifically modulating genetic risk genes are essential for development of disease-modifying treatments in AD [14]. A recent study showed that selecting genetically supported targets can double the success rate in clinical development [15].

However, existing data, including genomics, transcriptomics, proteomics, and interactomics (protein-protein interactions), have not yet been fully utilized and integrated to explore the roles of targeted therapeutic development for AD [16–18].

Understanding AD from the point-of-view of how human interactome perturbations underlie the disease is the essence of network medicine [13,14]. The main hypothesis of the AD network medicine is that cellular networks perturbed by genetic variants gradually rewire throughout disease pathogenesis and progression [13,14]. Systematic characterization and identification of underlying pathobiology will serve as a foundation for identifying disease-modifying targets for AD. Integration of the genome, transcriptome, proteome, and the human interactome are essential for such identification. In this study, we presented a <u>**net**</u>work <u>**t**</u>opology-based deep learning framework to identify disease-<u>**a**</u>ssociated <u>**g**</u>enes (NETTAG) and drug targets from genetic and genomic discoveries for AD. The key premises of NETTAG are that the disease risk genes: i) exhibit distinct functional characteristics compared to non-risk genes and therefore can be distinguished by their aggregated genomic features, ii) converge to a limited number of pathobiological pathways captured by the human protein-protein interactome, and iii) include multiple AD pathology modulators and potential therapeutic targets.

## Results

### A network-based deep learning framework

In this study, we presented NETTAG, a network-based deep learning framework to identify risk genes from GWAS and multi-omic findings in AD. NETTG integrates multi-omics data along with the human protein-protein interaction (PPI) network to infer likely risk genes and potential drug targets impacted by GWAS loci. Specifically, we assembled non-coding GWAS loci effects on expression quantitative trait loci (eQTLs), histone-QTLs, transcription factor binding-QTLs, enhancers and CpG islands, promoter regions, open chromatin, and promoter flanking region from GTEx [19], NIH RoadMap [20], Ensembl Regulatory Build [21], SNPnexus [22] and ENCODE [23] (**Figure 1**). The whole procedure is divided into 4 steps: i) We first utilized a deep learning model to cluster PPIs into multiple functional network modules by capturing its topological structures within the human protein-protein interactome (**Methods**). We then characterized each functional network module by linking its nodes (genes) with protein annotations from the Gene Ontology (GO) knowledgebase [24]; ii) We quantified node’s (gene’s) scores by integrating its functional similarity with each gene identified with multiple gene regulatory evidences via influencing GWAS loci; iii) We prioritized likely risk genes in AD by their aggregated gene regulatory features; and iv) we prioritized repurposable drugs for potential treatment of AD by evaluating network proximities between Alzheimer’s risk genes (alzRGs) and known drug targets under the human protein-protein interactome network model (**Figure 1**).

**Figure 1.**
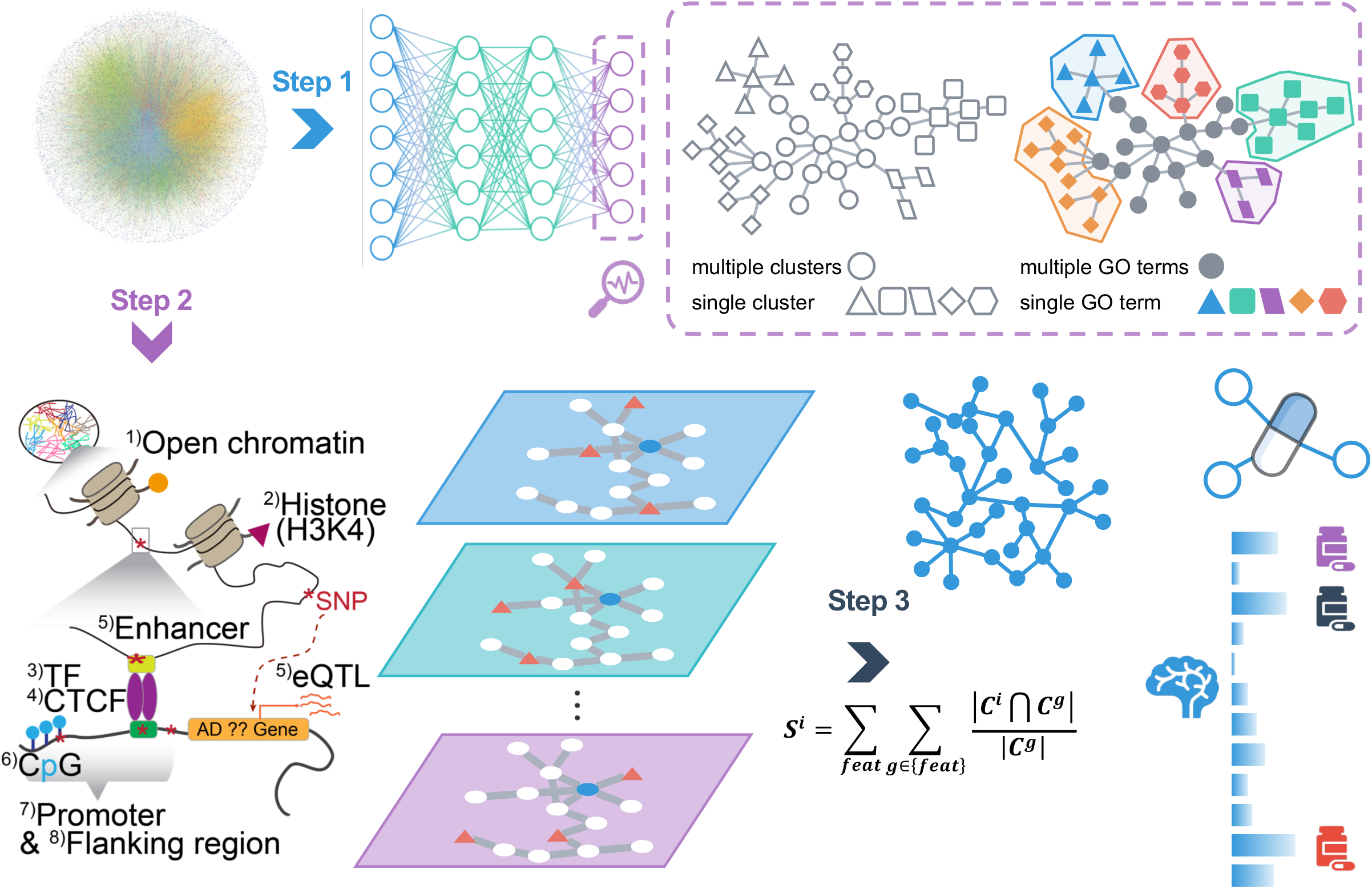
A diagram illustrating NETTAG. We first (Step 1) apply a deep-learning model to capture the topological structure of the PPIs and divide it into multiple subnetwork modules (**Methods**). Then we discover that the divided subnetwork module could approximate protein functions annotated by GO knowledge portal (**Methods**). Next (step 2), we predict AD-associated genes which are functionally similar as genes that have been identified by different gene regulatory elements, i.e., CpG island, CTCF, enhancer, eQTL, histone, open chromatin, promoter, promoter flanking region and transcriptional factors. Finally (Step 3), we prioritize repurposed drugs for potential treatment of AD by evaluating network proximities between predicted AD-associated genes and drug target networks (**Methods**).

### A gene regulatory landscape of GWAS loci in AD

After mapping AD loci (p<1.0×10^−5^) from multiple gene regulatory elements (**Methods**), we pinpointed 23 genes with CpG islands (e.g., *APOE*, *PVR*, *STK11*), 19 genes with CTCF binding sites (i.e., *BIN1*, *JPH1*, and *SYK*), 13 genes with enhancers (i.e., *BIN1*, *FARP1*, and *MARK4*), 21 genes with eQTL (i.e., *CD2AP*, *IL6*, and *PVR*), 169 genes with histone modifications including H3K27ac, H3K27me3, H3K36me3, H3K4me1, H3K4me2, H3K4me3, H3K9ac and H4K20me1 (such as *APOE*, *CKAP5*, *DST*, and *NECTIN2*), 48 genes with open chromatin (i.e., *BIN1*, *CLU* and *INPP5D*), 23 genes with promoter (i.e., *APOE*, *IL6*, and *STK11*), 59 genes with promoter flanking region (i.e., *BIN1*, *CLU* and *MARK4*), and 20 genes (i.e., *BCAM*, *CLU* and *VSNL1*) with transcriptional factor binding site, respectively (**Figure 2A**, **Supplemental Table S1**). As shown in **Figure 2A**, 69 genes have AD loci with multiple gene regulatory evidences, e.g. *APOE*, *BIN1*, *CLU*, *IL6*, *PTK2B* and *etc*. Specifically, *APOE* loci have regulatory evidences with CpG island (rs429358 and rs7412), histone (rs405509 and rs769449), promoter (rs769449), and promoter flanking regions (rs75627662) (**Supplemental Figure S1A**). Bridging integrator 1 (*BIN1*), another risk factor of LOAD [25][26], is associated with multiple regulatory elements as well, including CTCF binding sites (rs12989701), enhancer (rs10207628), histone (rs6431219, rs10194375, rs72838215, rs10207628), open chromatin (rs6733839), and promoter flanking region (rs10194375). Inositol polyphosphate-5-phosphatase D (*INPP5D*), a LOAD risk gene [10], is associated with open chromatin regulation (rs10933431). Mouse model (5XFAD) study found that expression of *Inpp5d* was elevated in microglia as the disease progressed [27]. Spleen associated tyrosine kinase (*SYK*) (rs1172922) is linked with the CTCF binding site. Activation of SYK boosts inflammation [28] and modulates both Aβ and tau-induced pathologies [29]. Phosphatidylinositol binding clathrin assembly protein (*PICALM*) loci in AD have multiple regulatory evidences, including histone (rs867611, rs527162, rs639012, and rs17817600), promoter (rs867611) and promoter flanking region (rs10792832). PICALM has been suggested to regulate AD pathology with Aβ generation and disorder lipid metabolism [30]. Altogether, these bioinformatics analyses highlight the crucial roles of gene regulation involving in various AD GWAS loci, which motivate us to develop NETTAG to infer new gene regulatory variants and putative risk genes in AD using network-based multi-omics evidence aggregation analyses.

**Figure 2.**
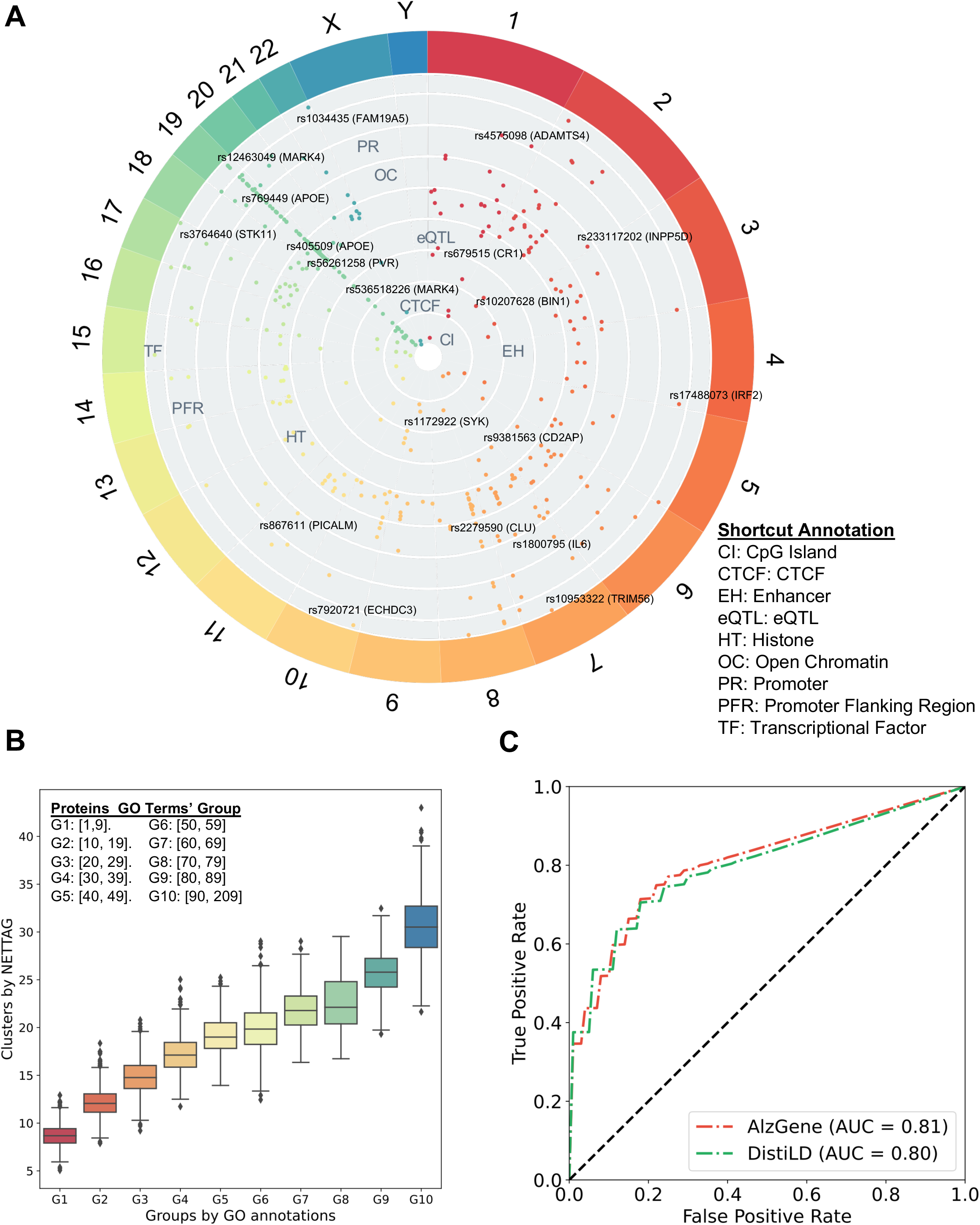
Gene regulatory landscape of AD GWAS loci. (*A*) Overview of AD GWAS loci across different chromosomes after considering nine gene regulatory elements: GpG island, CTCF, enhancer, eQTL, histone, open chromatin, promoter, promoter flanking region and transcriptional factors. (*B*) Proteins’ cluster numbers are positively correlated with their GO annotations. We divide proteins into 10 groups according to their GO terms. For example, G1 group include the proteins that have at least one, but less than ten GO annotation. (*C*) ROC analyses of NETTAG based on two collected AD-association gene sets, i.e., AlzGene and DistiLD.

### Network-based prediction of likely risk genes in AD

According to NETTAG, we first clustered PPIs into functional subnetwork modules using a deep learning framework. We found that the identified subnetwork modules could reflect biological relationships as well (**Figure 2B**). Specifically, proteins with more GO terms tend to have more clusters (**Figure 2B**, **Supplemental Table S2**). We found that proteins in the same subnetwork module tend to have more shared GO annotations (Wilcoxon signed rank test, p < 2.2×10^−16^, **Methods**). This indicates that network-based fingerprints of module overlays among genes can characterize functional modularities and similarities. We therefore inferred likely risk genes by integrating PPI-derived network modules and multimodal analyses of 9 types of gene regulatory impacts by AD GWAS loci. Specifically, taking CpG island as an example, the predicted score for one particular gene regarding CpG island could approximately estimate its functional overlap (spearman correlation, r = 0.44, p = 1.86×10^−24^, **Supplemental Figure S1B, Methods**) with all 23 AD CpG island-linked genes (**Supplemental Table S1**). Finally, we inferred likely risk genes by integrating (summing up) all 9 types of gene regulatory elements. The area under curve (AUC) for receiver operating characteristic curve (ROC) using AD-associated genes collected from AlzGene [31] and DistiLD [32] (**Figure 2C** and **Supplemental Figure 2A**) are 0.81 and 0.80 respectively, suggesting reasonable accuracy.

Via NETTAG (**Figure 1**), we identified 156 likely Alzheimer’s risk genes (termed alzRGs), such as *APOE*, *APP*, *BIN1*, *FYN*, and *STK11*. Among 156 alzRGs, products (proteins) of 139 alzRGs formed the largest connected component within 294 PPIs (**Figure 3A**, **Supplemental Table S3**). Via gene and functional enrichment analyses, we found that the NETTAG-predicted alzRGs are significantly enriched by gene regulatory elements (**Figure 3B** and **Supplemental Figure S3A**) compared to the same number of randomly selected genes with the similar degree distribution in the human interactome network. We assembled AD-associated genes from the GWAS catalog [33], UK Biobank GWAS [34] and DisGeNET with published experimental evidences from animal models and human studies [35]. We found that alzRGs were significantly enriched in all 3 AD-associated gene sets: GWAS catalog (adjusted p-value [q] = 2.25×10^−7^), UK Biobank GWAS (q = 8.59×10^−3^), DisGeNET (q = 1.19×10^−8^, Fisher’s exact test, **Supplemental Table S3**). Pathway enrichment analyses [36] showed that alzRGs are significantly enriched in multiple immune pathways (**Supplemental Table S3** and **Figure S3B**), including B cell (q = 5.32×10^−4^), T cell receptor (q = 1.17×10^−2^), cytokine signaling pathways (IL-2: q = 6.99×10^−3^, IL-7: q = 1.43×10^−2^, IL-18: q = 1.65×10^−2^). In summary, NETTAG achieved reasonable accuracy (AUC=0.81) in predicting likely AD risk genes with diverse functional pathways, including key immune pathways. We next turned to perform multi-omics validation for NETTAG-predicted alzRGs, including single-cell/nuclei transcriptomics in disease-associated microglia (DAM) and astrocyte (DAA) from transgenic mouse and human brains with well-known AD neuropathology.

**Figure 3.**
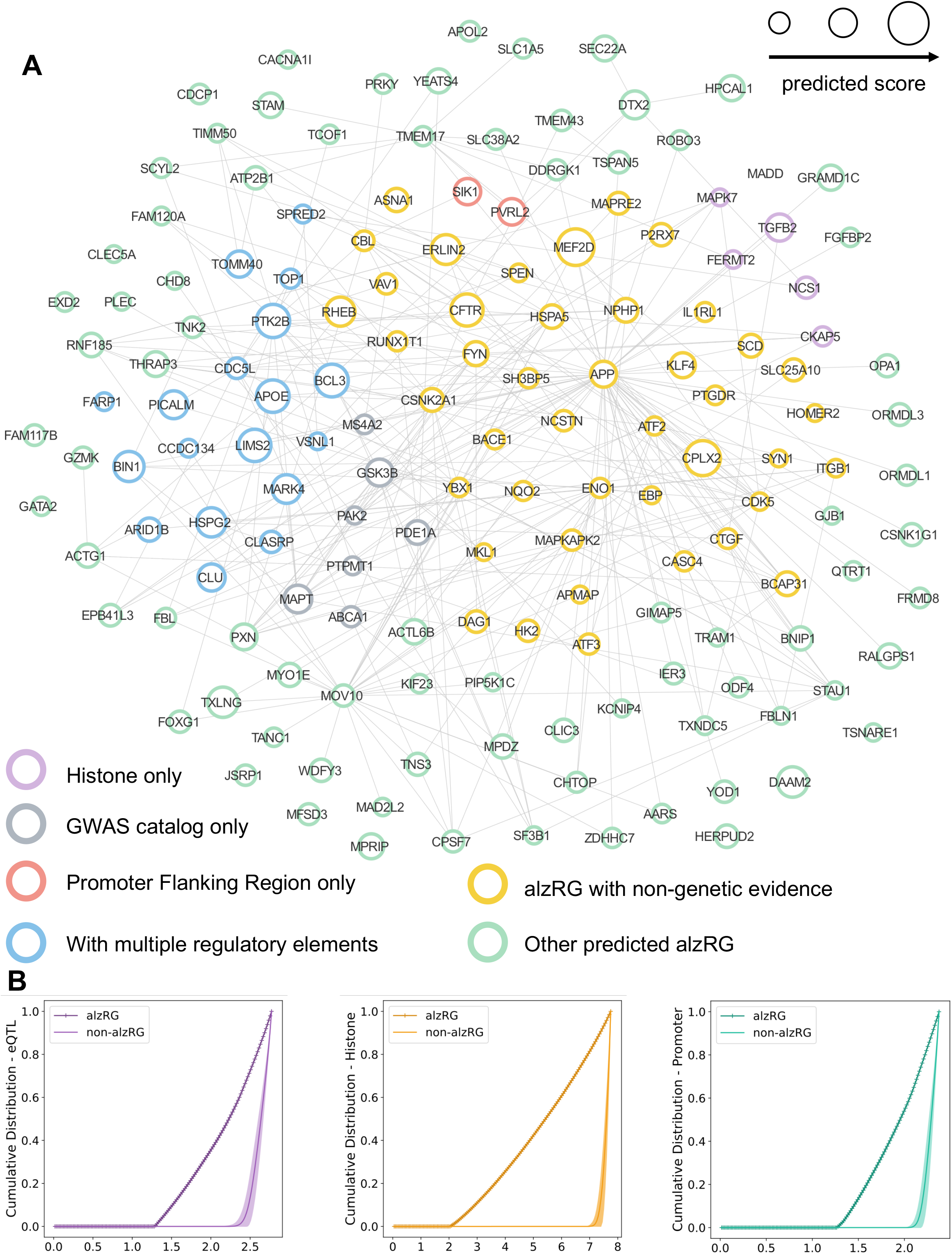
156 Prioritized AD-associated genes (alzRGs). (*A*) Visualization of 156 predicted alzRGs. And 139 alzRGs are non-isolated and form a subnetwork with 294 PPIs. Prioritized alzRGs are colored with various evidences. Purple circles denote genes that are exclusively related with histone regulatory element; red circles denote genes that are exclusively related with promoter flanking region regulatory element; blue circles denote genes that are simultaneously related with multiple gene regulatory elements; grey circles denote genes with no gene regulatory element associations but have been identified as AD-associated according to GWAS studies; yellow circles denote genes that are AD-associated by other types of evidences, e.g., animal models and etc.; green circles that predicted alzRGs with no previously reported evidences. (*B*) Cumulative distributions of predicted scores with alzRGs and same amount of random non-alzRGs with similar degree distribution for eQTL, histone and promoter gene regulatory elements, respectively.

### NETTAG-predicted genes are differentially expressed in AD

We found 95 alzRGs (p = 2.67×10^−7^, Fisher’s exact test) that are differentially expressed regarding at least one type of transcriptomics studies in AD. Specifically, 29 (p = 0.0185), 67 (p = 2.36×10^−3^), and 39 (p = 2.96×10^−7^) alzRGs are differently expressed genes (DEGs) according to microarray (human AD patients and controls), bulk RNA sequencing (human AD patients and controls), and single cell/nucleus RNA sequencing (AD transgenic mouse model and human postmortem brain samples) analyses, respectively (**Methods**). Nine genes (*ACTL6B*, *ATP2B1*, *EPB41L3*, *ABCA1*, *CPLX2*, *P2RX7*, *PDE1A*, *SLC38A2*, and *VSNL1*) are DEGs based on all three types of differential transcriptomic evidences (**Supplemental Figure S4A**). Block of purinergic receptor P2X 7 (P2RX7) was found to reduce tau accumulation in P301S tau transgenic mice [37]. Visinin like 1 (VSNL1) is co-expressed with multiple genes involving in molecular mechanisms of AD, including APP [38]. Nineteen genes (*ABCA1*, *APOE*, *BCL3*, *BIN1*, *CKAP5*, *CLU*, *FARP1*, *HSPG2*, *MADD*, *MAPK7*, *MARK4*, *NCS1*, *PICALM*, *PTK2B*, *SPRED2*, *TGFB2*, *TOMM40*, *TOP1*, and *VSNL1*) have been identified by gene regulatory elements and AD GWAS studies as well [33] (**Supplemental Figure S4B**). Microtubule affinity regulating kinase 4 (*MARK4*) is the one linked with most gene regulatory elements, including CpG island (rs28469095), CTCF (rs12463049), enhancer (rs536518226), eQTL (rs8100183), histone (rs9653111 and rs10421247), open chromatin (rs138137383), promoter flanking region (rs10421247 and rs138137383), TF-binding site (rs12463049) (**Supplemental Table S1**, **Supplemental Figure S4B**). MARK4 has been suggested as a potential target for AD via binding with acetylcholinesterase (AChE) inhibitors, such as donepezil (an AChE inhibitor used for patients with AD and other types of dementia) [39]. In addition to *APOE*, *BIN1* and *PICALM*, *CLU* loci are linked with four types of regulatory elements, including histone (rs9331896 and rs1532278), open chromatin (rs2279590), promoter flanking region (rs9331888), and TF-binding sites (rs1532278) (**Supplemental Table S1**, **Supplemental Figure S4B**). Clusterin (*CLU*) is another risk gene for LOAD [40] by involving in multiple AD pathologies, including neuroinflammation, Aβ accumulation, and lipid metabolism [41].

Among the 67 alzRGs that are differentially expressed based on bulk RNA-seq studies, 42 alzRGs form a subnetwork within human interactome network (**Figure 4A**). And within the 23 top differentially expressed alzRGs such as HOMER2, ABCA1, HSPG2 and etc. (|log_2_FC| > 1, q < 0.05, **Supplemental Table S4**), one downregulated gene homer scaffold protein 2 (*HOMER2*) (log_2_FC = −1.65, q = 2.39×10^−3^, **Supplemental Table S4**) was found to inhibit APP production and secretion of Aβ peptide together with HOMER3 [42]. ATP binding cassette subfamily A member 1 (*ABCA1*) was one top upregulated gene according to bulk RNA-seq studies (log_2_FC = 1.21, q = 1.37×10^−3^, **Supplemental Table S4**), and its mutation was associated with an elevated risk of AD [43]. Heparan sulfate proteoglycan 2 (*HSPG2*) was another top upregulated gene according to bulk RNA-seq studies (**Supplemental Table S4**), it was suggested that people carrying APOE epsilon4 allele have increased risk of AD if carrying HSPG2 A allele at the same time [44].

**Figure 4.**
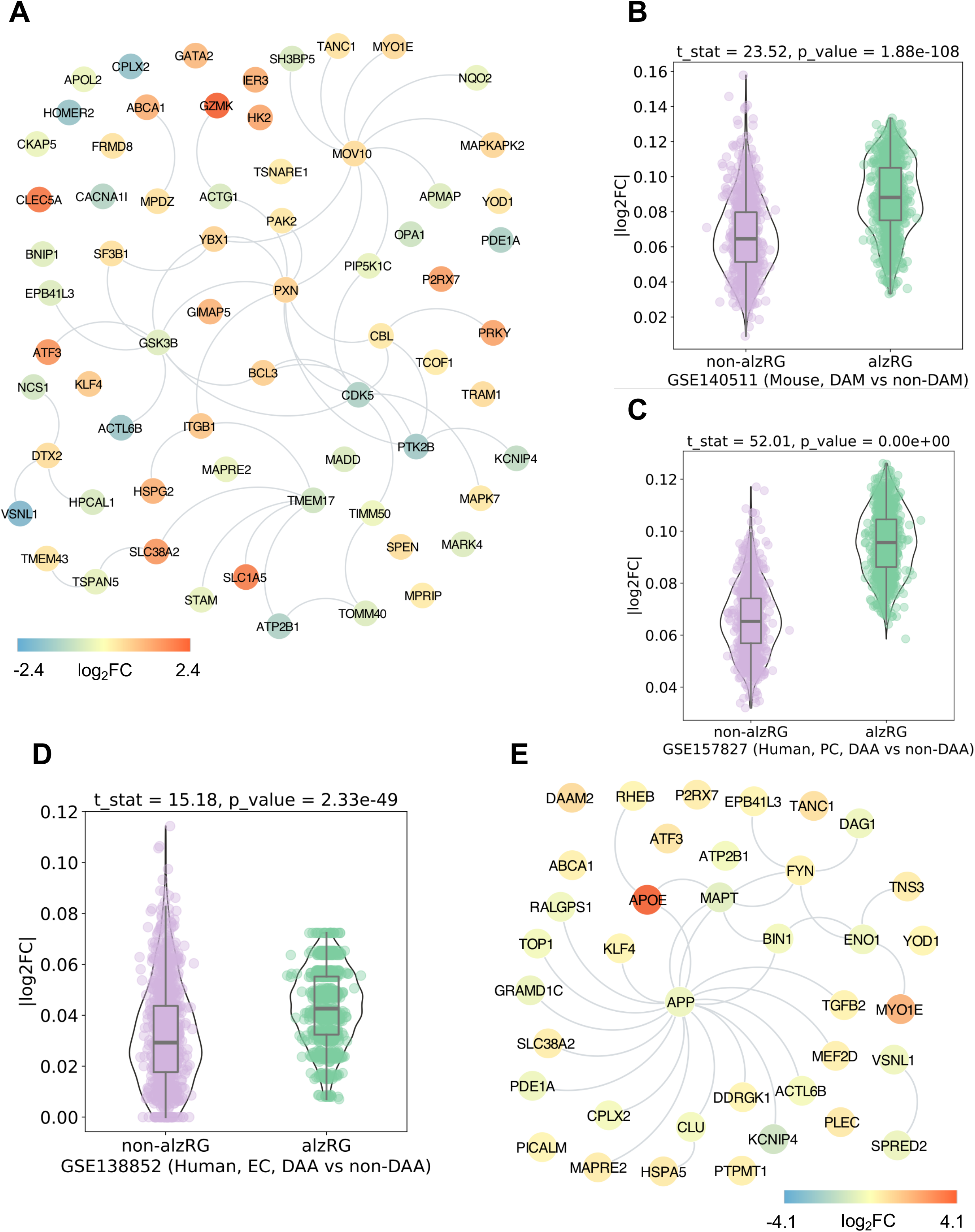
Predicted alzRGs are more likely differently expressed according to transcriptome studies (*A*) Visualization of 67 predicted alzRGs that are also DEGs according to human bulk RNA-seq studies with both LAD and control donors (**Methods**). (*B*) Violin plots show alzRGs are more likely differentially expressed in DAM according to mouse single-nucleus (GSE140511) RNA seq datasets. (*C*) Violin plot shows alzRGs are more likely differentially expressed in DAA according to human prefrontal cortex (PC) single-nucleus RNA seq dataset (GSE157827). (*D*) Violin plot shows alzRGs are more likely differentially expressed in DAA according to human entorhinal cortex (EC) single-nucleus RNA seq dataset (GSE157827). (*E*) Visualization of 32 predicted alzRGs that are also DEGs according to sc/sn RNA-seq studies collected from both mouse models and human postmortem brain tissues in two disease-associated immune subtypes, i.e., DAM and DAA (**Methods**).

We have collected 6 sets of proteomics data from transgenic mouse model (**Methods**). Among the 156 predicted AD-associated genes, protein products of 29 genes (i.e., *HSPA5*, *ENO1*, *FERMT2* and *VAV1*) are differentially expressed (p = 3.98×10^−7^, Fisher’s exact test, **Supplemental Table S4**). Heat shock protein family A (Hsp70) member 5 (*HSPA5*) was suggested with important roles in tau phosphorylation and a potential target for AD treatment [45]. Study showed that oxidative inactivation of enolase 1 (ENO1) could accelerate development of AD from mild cognitive impairment (MCI) [46]. FERM domain containing kindlin 2 (*FERMT2*) is identified as AD risk genes by GWAS studies [10]. It is also suggested that FERMT2 could modulate APP metabolism and Aβ formation, therefore linking its mechanism association with AD [47]. Another study in mouse model found that targeting vav guanine nucleotide exchange factor 1 (VAV1) could rescue neuron death by inhibiting JNK signaling pathway [48].

### alzRGs are differentially expressed in AD-associated microglia and astrocytes

Neuroinflammation plays crucial roles in pathogenesis and disease progression of AD [49]. We found that predicted alzRGs are significantly enriched by multiple immune pathways, including B cell receptor, T cell receptor, and cytokines (IL-2, IL-7 and IL18) signaling pathways (**Supplemental Table S3**, **Supplemental Figure S3B**). We next turned to investigate how inflammatory pathways were impacted by predicted alzRGs using disease-associated microglia (DAM) and disease-associated astrocytes (DAA) as examples. We found that 14 alzRGs were differentially expressed (|log_2_FC| > 0.25, q < 0.05) in DAM from one 5XFAD mouse model derived single-cell RNA-seq dataset (onesided t-test: statistic = 33.85, p = 9.76×10^−183^, **Supplemental Figure S4C**) and one 5XFAD mouse model derived single-nucleus RNA-seq dataset (one-sided t-test: statistic = 23.52, p = 1.88×10^−108^, **Figure 4B**). For DAA, 25 genes are differentially expressed (|log_2_FC| > 0.25, q < 0.05) across 3 snRNA-seq datasets from human postmortem brains with different brain regions, including prefrontal cortex (p < 1.0×10^−3^, **Figure 4C**), entorhinal cortex (p = 2.33×10^−49^, **Figure 4D**), and super frontal gyrus (p = 8.30×10^−27^) (**Supplemental Figure S4D**, **Supplemental Table S4, Methods**). Among 39 differentially expressed alzRGs in DAM or DAA, 28 alzRGs form a subnetwork with human interactome network (**Figure 4E**). Activating transcriptional factor 3 (*ATF3*) overexpressed in DAM (one top DEGs log2FC = 0.70, q = 2.21×10^−21^ with 5XFAD mouse models, **Supplemental Table S4**), was also observed with elevated expression in another mouse model study [50]. The study also suggested that the elevated expression level of ATF3 is positively correlated with Aβ accumulation [50]. Microtubule associated protein RP/EB family member 2 (MAPRE2) overexpressed in DAA (one top DEGs log2FC = 0.51, q = 7.03×10^−20^ with human postmortem brain tissues, **Supplemental Table S4**) was identified as a new AD-associated gene based on GWAS from ADNI cohort [51].

### Discovery of high-confidence risk genes in AD

We next turned to identify high-confidence risk genes in AD via combining multiple factors: 1) high-confidence risk genes are top predicted by NETTAG (156 alzRGs); 2) high-confidence risk genes are supported with at least 3 types of multi-omics evidences (**Supplemental Table S4**); 3) high-confidence risk genes have not previously been identified by GWAS Catalog. In total we have identified 25 high-confidence AD risk genes, e.g., *BACE1*, *CDK5*, *CPLX2*, *FYN*, *MAPKAPK2*, *MEF2D*, and etc. (**Figure 5, Supplemental Table S4**). Myocyte enhancer factor 2D (*MEF2D*), an alzRG with currently non-existing AD-associated DNA regulatory or GWAS evidence, is the one with the highest predicted score. It is differently expressed regarding to both singlenucleus RNA seq and microarray analyses (**Figure 5, Supplemental Figure S4E, Supplemental Table S4**). One study found that protocatechuic acid could rescue a cell model from okadaic acid-induced cytotoxicity (tau hyperphosphorylation) by modulating Akt/GSK-3β/MEF2D pathway and exhibit neuroprotective effects which may suggest itself being beneficial for AD treatment [52]. Complexin 2 (*CPLX2*) is our second top predicted alzRG with currently non-existing AD-associated DNA regulatory or GWAS evidence. It is identified as AD-associated with five types of evidence and differently expressed according to both transcriptome (DAA) and proteome studies (**Figures 5, Supplemental Figure S4F**, **Supplemental Table S4**). Experiment with hippocampus from 3x-Tg AD mice showed abnormal lower proteomic levels of CPLX1 and CPLX2 [53] which is consistent with the transcriptomic behavior in DAA. Further decreased expression level was observed after exposure to copper, which suggested that CPLX2 together with CPLX1 may be the key factors in chronic copper overexposure-induced memory damage in AD [53].

**Figure 5.**
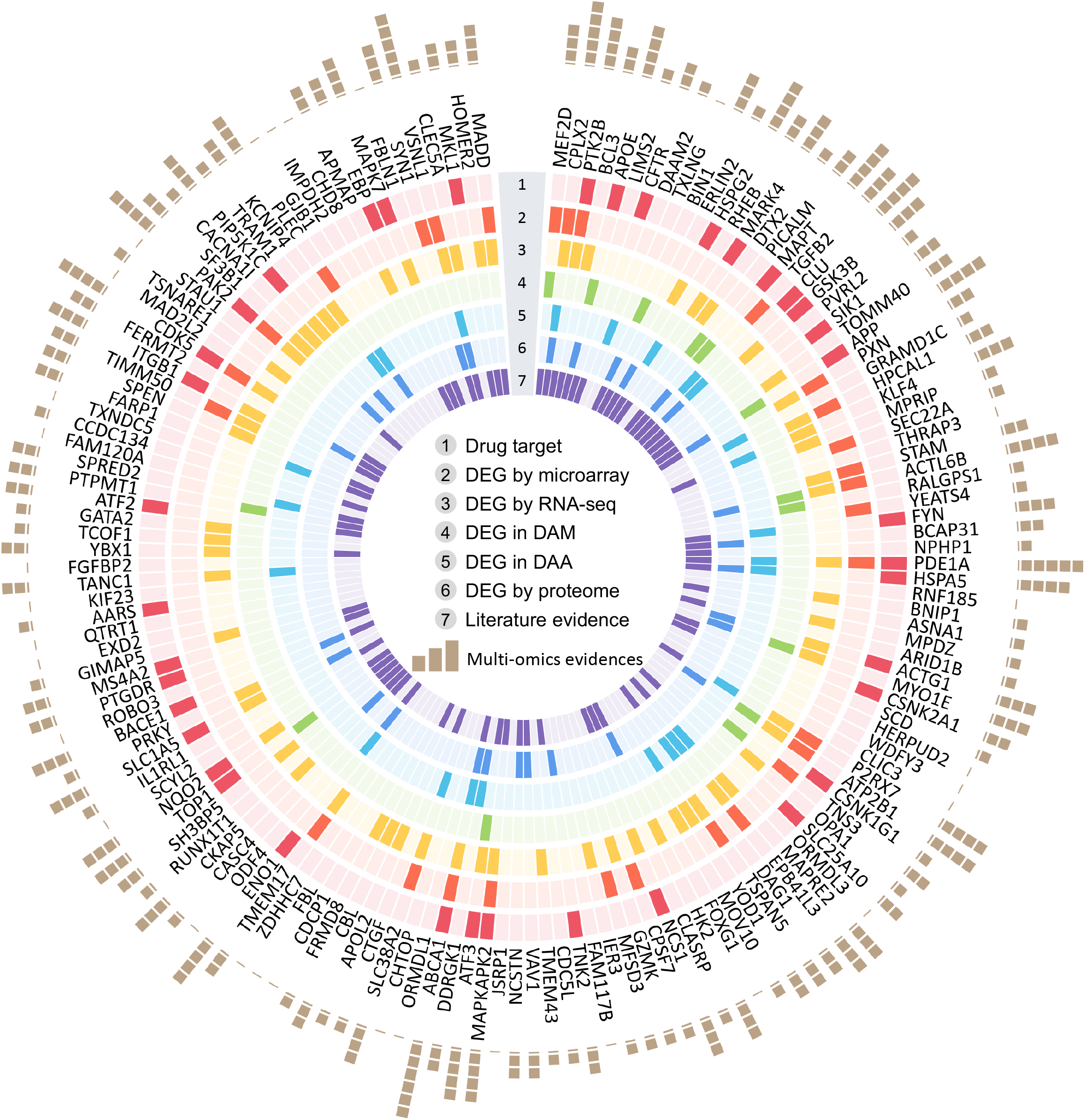
Predicted high-confidence AD-associated genes (*A*) Summary of multi-omics validations for all 156 predicted alzRGs with seven types of evidences. The genes are sorted in predicted score descreasing order (clockwise direction). We have collected seven types of evidences, including drug target, DEG by microarray studies, DEG by bulk RNA-seq studies, DEG in disease-associated microglia (DAM), DEG in disease-associated astrocyte (DAA), DEG by proteome studies and literature evidences. There are 126 predicted alzRGs that could be proved as AD-associated with at least one type of evidence.

### Discovery of repurposable drugs via targeting risk genes

Among the 156 predicted alzRGs, 38 proteins have been identified as known drug targets (p = 8.78×10^−4^, Fisher’s exact test, **Supplemental Table S4** and **Figure 5**). In total, 9 targets (i.e., APP, ATF2, BACE1, CDK5, FYN, GSK3B, MARK4, MKL1 and PTK2B, **Supplemental Table S4**) have been widely investigated as therapeutic approaches for treating AD. FYN proto-oncogene, Src family tyrosine kinase (FYN), which contributes to Aβ production and tau phosphorylation, has been suggested as one potential target for AD [54]. Glycogen synthase kinase 3 beta (GSK3B) has been found in hyperphosphorylation of tau and Aβ production [55]. Study has found that one GSK3B inhibitor thiadiazolidinone could decrease tau phosphorylation and improve neuronal survival [55]. Beta-secretase 1 (BACE1), a β-secretase enzyme involving in Aβ peptide generation, has been demonstrated a promising target in AD [56]. We next turned to identify repurposable drugs by specifically targeting alzRGs (**Figure 3A**).

Using our well-established network proximity approaches [57], we computationally identified 118 candidate drugs using z-score < −2 and q < 0.05 from total 2,938 U.S. FDA-approved or clinically investigational drugs (**Supplemental Table S5, Methods**). As shown in **Figure 6A**, we grouped these top predicted 118 candidate drugs into 14 pharmacological categories based the first-level of the Anatomical Therapeutic Chemical (ATC) code. Choline, a nutrient found in many vitamins, is our 5^th^ ranked predicted drug (**Supplemental Table S5**). Experiments with APP/PS1 mouse models showed that dietary of choline could reduce Aβ production and improve spatial memory by suppressing overactivation of DAM [58]. Choline does not directly target any alzRGs; yet, choline’s targets interacts with multiple protein products of predicted alzRGs, including APP, BIN1, CDK5, and FYN (**Figure 6B**). Ibudilast, an anti-inflammatory drug used to attenuate multiple sclerosis, is another top prediction (**Supplemental Table S5**). Ibudilast inhibited pro-inflammatory cytokine production and blocked neuroinflammation to prevent Aβ-produced cognitive impairment [59]. Mechanistically, ibudilast’s targets (i.e., PDE3A, PDE4B and PDE4D) have physical interactions with proteins encoded by several predicted alzRGs (i.e., BIN1, FYN, GSK3B) (**Figure 6C**). In summary, risk genes identified by NETTAG offer potential drug targets for AD therapeutic discovery, including drug repurposing (such as ibudilast and choline). The predicted repurposable drugs offer potential candidates for future preclinical and clinical validations. Deferoxamine, a FDA-approved iron-chelating agent for treatment of iron overdose or hemochromatosis, is the top second predicted candidate drug (**Supplemental Table S5**). Treatment with deferoxamine alleviated disturbed iron homeostasis and reduced APP phosphorylation in a transgenic mouse model [60]. Mechanistically, several deferoxamine’s targets (HIF1A, RGS4, RGS19 and GMNN) have physical protein interactions with multiple protein products of several predicted alzRGs, such as APOE, BIN1, CLU and GSK3B (**Supplemental Figure S4G**).

**Figure 6.**
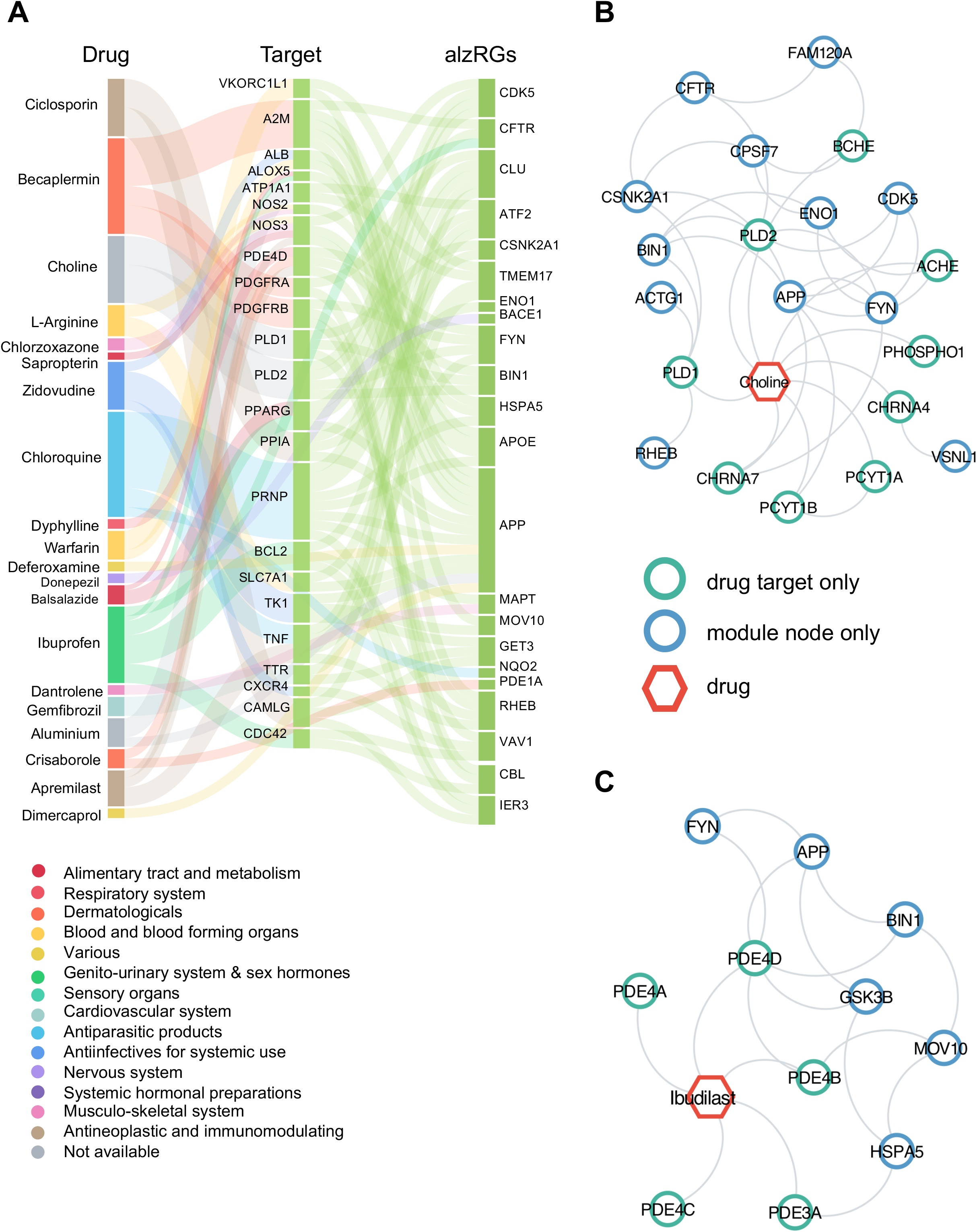
Network-based discovery of repurposable drug candidates for AD by evaluating the network proximity between predicted alzRGs and drug-target networks (*A*)118 prioritized drugs for AD treatment. Drug are grouped by fourteen different classes (e.g., immunological, respiratory, neurological, cardiovascular, and cancer (**Supplemental Table S5**)) defined by the first-level of the Anatomical Therapeutic Chemical (ATC) codes. (*B*) Proposed mechanism-of-action (MOAs) for Choline by drugtarget network analysis. (*C*) Proposed MOAs for Ibudilast by drug-target network analysis.

## Discussion

In this study, we presented a deep-learning framework that integrates multi-omics knowledges to infer putative risk genes in AD. To avoid “black-box” deep learning models, we utilized human protein-protein interactome network to make NETTAG more transparent and interpretable in human gene discovery. For example, (**Methods**), there are 16,720 proteins in the used human protein interactome according to the GO knowledgebase [24] (**Supplemental Table S2**). We found that 97% (16,214) of annotated proteins in the human interactome have multiple GO terms, ranging from 2 to around 200 (**Supplemental Table S2**, **Supplemental Figure S5A**). We further found that the deep learning-predicted subnetwork modules were highly correlated with protein functions (**Figure 2B**). We performed additional spearman (r) correlation analysis to evaluate the correlation between predicted score (cumulative overlays of divided subnetwork modules) and cumulative overlays of protein functions by considering each gene regulatory element. We found that deep learning-predicted scores showed significant correlations across all gene regulatory elements (**Supplemental Figure S1B**). For example, CpG island (r = 0.44, p = 1.86×10^−24^, enhancer (r = 0.34, p = 9.07×10^−13^), histone (0.50, p = 6.49×10^−37^), and TF-binding site (0.36, p = 3.50×10^−17^), show strong correlations; while promoter flanking region (0.26, p = 2.60×10^−12^), open chromatin (0.22, p = 2.91×10^−10^), promoter (0.20, p = 2.42×10^−15^), CTCF (0.09, p =1.45×10^−2^) and eQTL (0.14, p = 8.62×10^−10^) show weak correlations. These observations showed that CpG island, enhancer, histone and TF-binding sites play more important gene regulatory roles of GWAS loci in AD compared to promoter flanking region, open chromatin, promoter, CTCF and eQTL.

We performed ROC analyses to evaluate the performance of NETTAG. We found that the AUCs for predicted scores by integrating nine DNA regulatory elements are 0.23 (AlzGene) and 0.23 (DistiLD) higher than the averages of AUCs if considering each single regular element alone (**Supplemental Figures S2B-2D**). This suggested integrating multi-genomic evidences could infer like risk genes with a higher accuracy. We acknowledged several potential limitations in the current study. For example, incompleteness of the human protein-protein interactome and GWAS loci by lack of population size may influence the model performance. In addition, the genome regulatory elements used in this study is not brain-specific. More brain-specific functional genomics data should be integrated in the near future [61]. If considering all protein pairs sharing at least one common GO annotations, we found that shortest path-based distances among those protein pairs are 2 (53.97%), 3 (39.05%) and 4 (5.05%) in decreasing order (**Supplemental Figure S5B**). Therefore, in NETTAG, we have 2 hidden layers to indirectly aggregate features from second order neighbors. We postulate that reformulating the model which could sampling second and third order neighbors directly may improve the model performance further.

In summary, we established a deep learning framework (NETTAG) that incorporates multi-genomics knowledges along with human PPIs to infer novel AD-associated genes. We showed that predicted genes are enriched with drug targets, differently expressed in disease-associated immune cell subtypes, and most importantly significantly AD-associated. We believe that the NETTAG presented here, if broadly applied, would significantly catalyze innovation in AD drug discovery.

## Methods and Materials

### Construction of genetic features

In this study, we collected 1,047,489 SNPs across multiple genetic traits from GWAS catalog [33], such as Alzheimer’s disease, cerebral amyloid deposition measurement. Next, we performed web server SNPnexus [22] to annotate all SNPs in human genome (GRCh38) and collected the regulatory elements information from five databases, including GpG Islands [22], Ensembl Regulatory Build [21], ENCODE [23], the Genotype-Tissue Expression (GTEx) portal [19] and Roadmap [20]. Finally, nine regulatory elements (histone, open chromatin, CpG Island, TF, CTCF, eQTL, enhancer, promoter and promoter flanking region) were used as features to evaluate AD disease-associated genes. To be more specific, <u>**step 1**</u>: for SNPs with respect to each regulatory elements, e.g., CpG island, we merge them with SNPs curated by GWAS Catalog with AD as the mapped traits. <u>**Step 2**</u>: For each AD related SNPs, the corresponding genes were identified by the “MAPPED GENE(S)” column as provided by GWAS Catalog. For SNPs with no mapped genes, if there were any reported genes (REPORTED GENE(S) column in GWAS Catalog) associated with this SNP, we then map the SNP to its reported genes. For ENCODE [23] and Ensembl Regulatory Build [21] databases, we only consider epigenomes from brain and neuron tissues and normal karyotype. The specific epigenomes with brain and neurons for each database are presented in **Supplemental Table S1**, separately. The final mapped genes for each regulatory element are provided in **Supplemental Table S1**.

### Building Human Protein-protein interactome

To build the comprehensive human interactome from the most contemporary data available, we collect 18 commonly used PPI databases with experimental evidence which mainly include: (i) binary PPIs tested by high-throughput yeast-two-hybrid (Y2H) systems [16]; (ii) kinase-substrate interactions; (iii) signaling networks; (iv) binary PPIs from three-dimensional protein structures; (v) protein complexes data; and (vi) carefully literature-curated PPIs. In total, 351,444 PPIs connecting 17,706 unique proteins are free available at https://alzgps.lerner.ccf.org. And in this study, we only consider its largest connected components which includes 17,456 proteins and 336,549 PPIs. All details are provided in our recent studies [62,63].

### Description of NETTAG

NETTAG involves 3 steps. <u>**Step 1**</u>: we build up a graph neural network (GNN) model to capture PPI’s topology structure and establish the appropriate overlapping clustering. The GNN model is motivated by NOCD-G [64] which is one GNN-based overlapping community detection framework. And in NETTAG, we made 2 modifications with respect to NOCD-G [64].

Modification 1: The model architecture used in NETTAG is defined below (Eq.1):

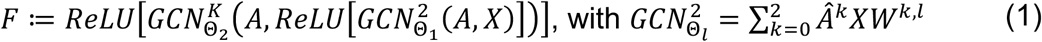

Here: 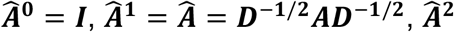 is the elementwise square of 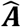 and the sub/super-script *l* is the layer index. *A* is the adjacency matrix of the PPI, *D* is the corresponding diagonal degree matrix. *X* in generate denotes the node feature matrix, and here we set *X* = *A* as implemented by NOCD-G [64]. In classical graph convolution network (GCN) models [65], nodes with various degrees share the same weight matrix. Therefore in NETTAG, coupling 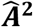 into the model is to overcome this, i.e., to assign different weight matrices for nodes with high and low degrees. To evaluate its effects, we compare the predicted genes generated by NETTAG with (test group) and without (control group) this extra 2nd order term. Each group has 10 experiments with fixed seeds. We find that predicted alzRGs generated by models from test group are more significantly AD-associated compared with those from the control group (paired seeds) with multiple sources, i.e., GWAS catalog (q without / with 2nd order term: 3.07×10^−3^ ± 7.92×10^−3^ / 3.53×10^−5^ ± 1.07×10^−4^, **Supplemental Figures S5C**), UK Biobank GWAS (q without / with 2nd order term: 1.22×10^−2^ ± 1.53×10^−2^ / 7.37×10^−3^ ± 3.92×10^−3^, **Supplemental Figures S5D**), and DisGeNET (q without / with 2nd order term: 6.30×10^−5^ ± 1.43×10^−4^ / 8.65×10^−7^ ± 2.67×10^−6^, **Supplemental Figures S5E**). Finally, the dimension of the final output layer interprets the clustering number.

The output matrix *F* is then feed into Bernoulli-Poisson model [66] (**Eq.2.1**) to learn PPI’s topology. The output matrix *F* has N (number of total nodes in PPI) rows and C (clustering numbers) columns. Each specific row (vector) denotes the node’s weights for being assigned to each cluster. Therefore, we can interpret the loss as follows: if two nodes have multiple commonly shared clusters (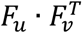 is large), then there should exist an edge connecting each other (1 - exp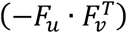 is close to 1) and vice versa.

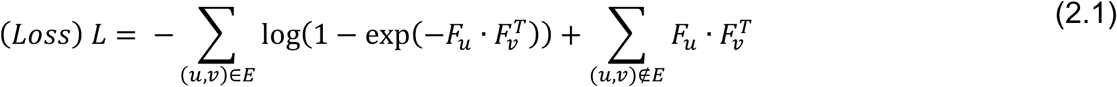

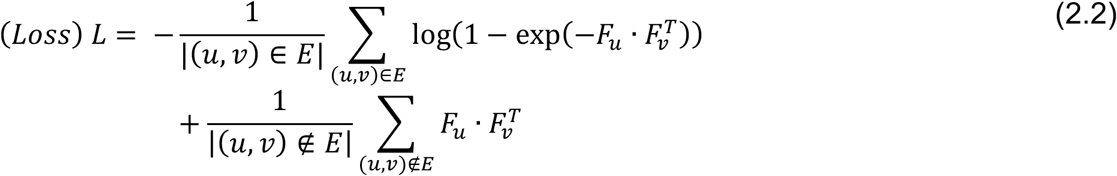

<u>Modification 2:</u> The PPI network is a highly sparse network which implies number of connected edges is far less than that of non-connected edges (For our PPI, the ratio between number of connected edges over number of non-connected edges = 1 / 448). To address this imbalanced training problem, the authors in [64] first uniformly subsampled certain amounts of connected edges and then subsampled the equal amounts of non-connected edges to train the averaged loss (**Eq 2.2**). Instead of uniformly sampling edges directly, we first group nodes according to their degrees into multiple bins. And in each training iteration, we first uniformly subsample same amounts of nodes from each bin, then we extract an adjacency matrix *A_sub_* which compromising only selected nodes. In this way, we can keep the topology similarities among sampled subgraphs from different iterations. Next, we use the connected and non-connected edges in *A_sub_* to compute the train loss, and the rest connected and non-connected edges in *A*\*A_sub_* for test loss evaluation. With this graph subsampling scheme, we find that we are capable to maintain similar connected and non-connected training edge percentages (**Supplemental Figure S5F).**

After learning the clustering affinity matrix *F*, we used the threshold defined in [67] as the cutoff for *F* to determine the node (gene) clustering membership (**Eq. 3**).

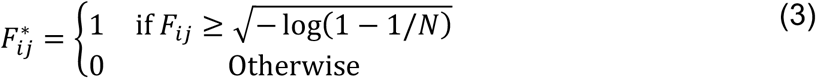

#### Step 2 of NETTAG

After clustering the whole PPI into overlapping communities, we next score each node with respect to each gene regulatory element. The detailed steps are explained in **Supplemental Materials**. Each node (gene) is firstly scored according to its clustering overlap with regulatory element evidenced genes. Next, we construct the background distribution by performing 1,000 random experiments to evaluate the statistical significance of node score computed in previous step. Finally, we evaluate the integrated node score by summing up all scores with respect to each gene regulatory element.

#### Step 3 of NETTAG

After inferring the nodes’ score, finally we extract the disease module by considering genes which are inferred only significantly associated with AD. In detail, we collect all positive gene scores {S_*i*_ > 0}, and compute the mean *μ*, standard deviation *σ* and the z score. We consider genes with z-scores great than 2.32 (p value = 0.01) as the predicted AD associated genes, and map those genes to the background PPI network to generate disease module.

### Network proximity for drug prediction

We assembled drugs from the DrugBank database relating 2,938 compounds [68]. To predict drugs with extracted disease module from NETTAG, we adopted the closest-based network proximity measure [57] as below.

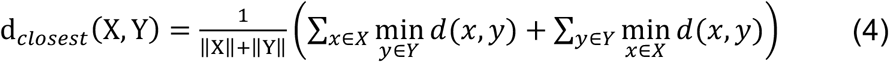

where d(x,y) is the shortest path length between gene x and y from gene sets X and Y, respectively. In our work, X denotes the disease module from NETTAG, Y denotes the drug targets (gene set) for each compound. To evaluate whether such proximity was significant, the computed network proximity is transferred into z score form as shown below:

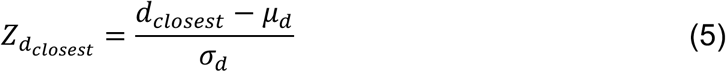

Here, μ_d_ and σ_d_ are the mean and standard deviation of permutation test with 1,000 random experiments. In each random experiment, two random subnetworks X_r_ and Y_r_ are constructed with the same numbers of nodes and degree distribution as the given 2 subnetworks X and Y, separately, in the PPI network.

### Agreement between clustered subnetwork modules and protein functions

With respect to each clustered subnetwork modules, considering its all-possible protein-protein pairs, we count numbers of protein-protein pairs that share at least one common GO annotations and numbers of protein-protein pairs that share no common GO annotations. Then we build up a paired statistical test with null hypothesis as proteins in the same clustered subnetwork modules have no protein functional similarities, and alternative hypothesis as proteins in the same clustered subnetwork module possess common protein functions. We apply Wilcoxon signed rank test with R.

### Differentially expressed gene (DEG) with transcriptome and proteome analyses

The transcriptome analyses are performed based on microarray, bulk RNA-seq, and single-cell/nucleus (sc/sn) RNA-seq datasets. We utilize three sets of human brain microarray transcriptome data collected from late-stage AD and control donors. They are available from Gene Expression Omnibus (https://www.ncbi.nlm.nih.gov/geo/) database under accession numbers: GSE29378 (31 LAD and 32 controls) [69], GSE48350 (42 AD and 173 controls) [70], and GSE84422 (328 LAD and 214 controls) [71]. We also include human brain bulk RNA-seq transcriptome data collected from hippocampus region of late-stage AD and control donors with three studies including 4 LAD versus 4 controls [72], 6 LAD versus 6 controls [73], and 20 LAD versus 10 controls [74]. The DEGs for all microarray and bulk RNA-seq datasets have been analyzed in recent developed AD knowledgebase AlzGPS [75]. The complete sc/sn RNA-seq datasets used for DEG analyses in this study are available from Gene Expression Omnibus (https://www.ncbi.nlm.nih.gov/geo/) database under accession numbers: GSE98969 [76], GSE140511 [77], GSE147528 [78], GSE138852 [79] and

GSE157827 [80]. We have performed the corresponding differently expressed gene analyses in our previous study [63] for datasets GSE98969, GSE140511, GSE147528 and GSE138852. The more detailed bioinformatics analysis for newly add dataset GSE157827 is described in **Supplemental Materials**. The DEG analysis between DAM and non-DAM is based on mouse sc-RNA seq datasets GSE98969 and mouse sn-RNA seq dataset GSE140511. The union of DEGs between DAM and homeostasis microglia from both datasets are used for transcriptome analysis in this study. The DEG analysis between DAA and non-DAA is based on the rest three human sn-RNA seq datasets GSE147528, GSE138852 and GSE157827. The union of DEGs between DAA and homeostasis astrocytes from all these three human datasets are used for transcriptome analysis in this study. For all DEGs generated from sc/sn RNA-seq datasets, we apply uniform criterion with q < 0.05 and |log_2_FC| ≥ 0.25. Proteome analyses are based on six mouse model datasets including 7- and 10-months ADLP mouse models (JNPL3 mouse model cross with 5XFAD mouse model) [81], 7- and 10-months 5XFAD mouse models [81], 12 months 5XFAD mouse model [82] and 12 months hAPP mouse model [82].

### Collections of AD seed genes

We collect AD seed genes from two AD-included knowledgebase, including AlzGene and DistiLD. AlzGene [31] collected AD-associated genes via genetic association studies. Thirty-two genes supported by genetic evidences are collected from AlzGene. DistiLD made the existing GWAS studies easier for accessing disease-associated SNPs and genes [32]. We collected 19 genes (p < 5.0×10^−8^) with AD GWAS from DistiLD. The complete lists of genes we used for ROC analyses are presented in **Supplemental Table S2**.

### Enrichment Analysis

All pathway and disease enrichment analyses were conducted using WikiPathways [83], GWAS Catalog 2019 [33], UK Biobank GWAS v1 [34] and DisGeNET [35] from Enrichr [84], respectively.

### Gene Ontology

All proteins’ gene ontology annotations (human, gaf-version 2.1) are extracted from The Gene Ontology (GO) Knowledgebase [24]. For our PPI with 17,706 proteins, there are 16,736 proteins with total 268,241 GO annotations.

## Software availability

All codes written for and used in this study are available from https://github.com/ChengF-Lab/NETTAG.

## Supplementary information

is available in the online version of the paper.

## Author Contributions

F.C. conceived the study. J.X. performed all experiments and data analysis. Y.H. performed multi-omics data analyses. Y.Z. and M.H. interpreted the data analysis. J.X. and F.C. drafted the manuscript and critically revised the manuscript. All authors critically revised and gave final approval of the manuscript.

