## Supplementary Figures for "A Network-based Deep Learning Framework Catalyzes GWAS and Multi-Omics Findings to Biology and Drug Repurposing for Alzheimer’s Disease"

### Supplemental Figure S1

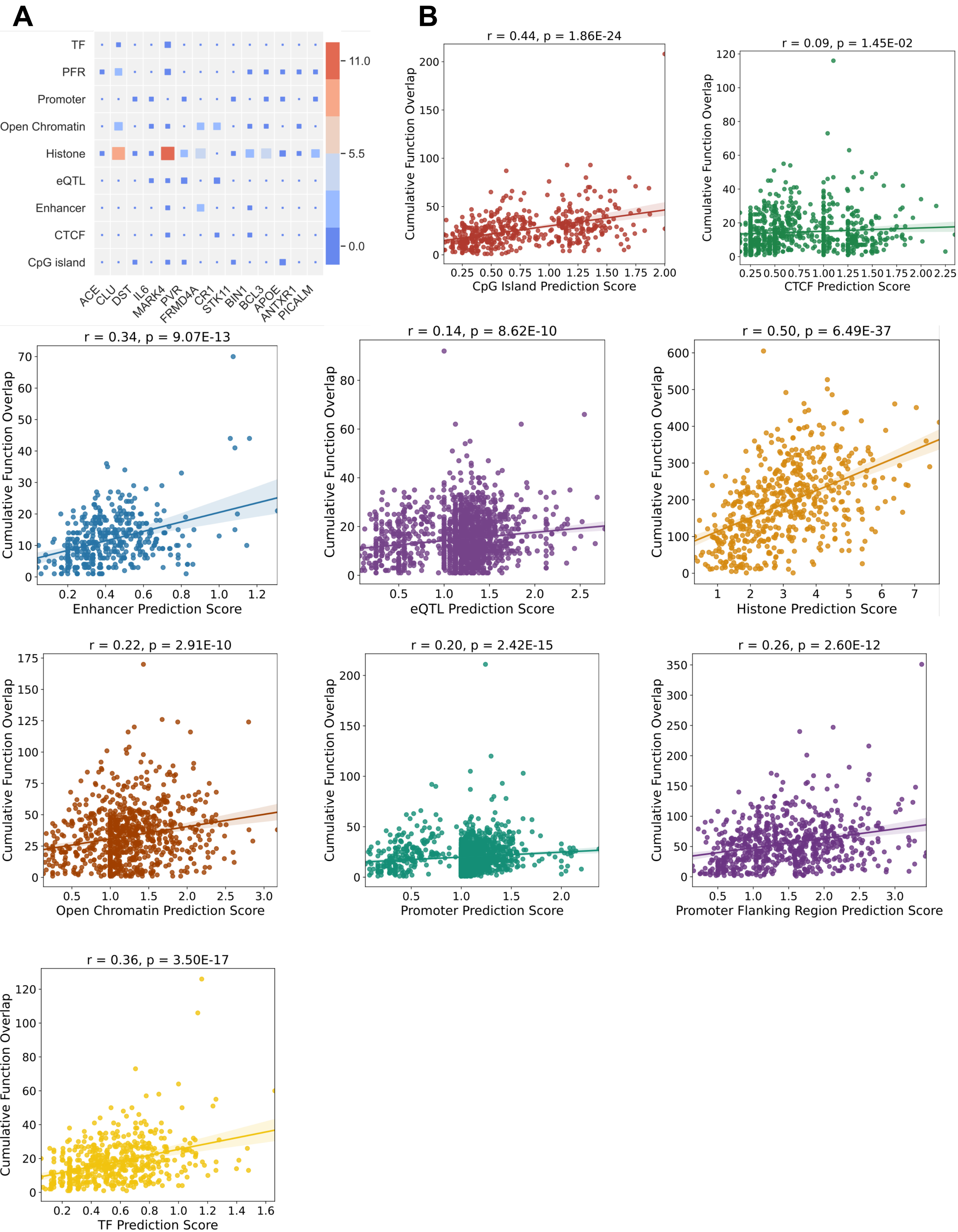

### Supplemental Figure S2

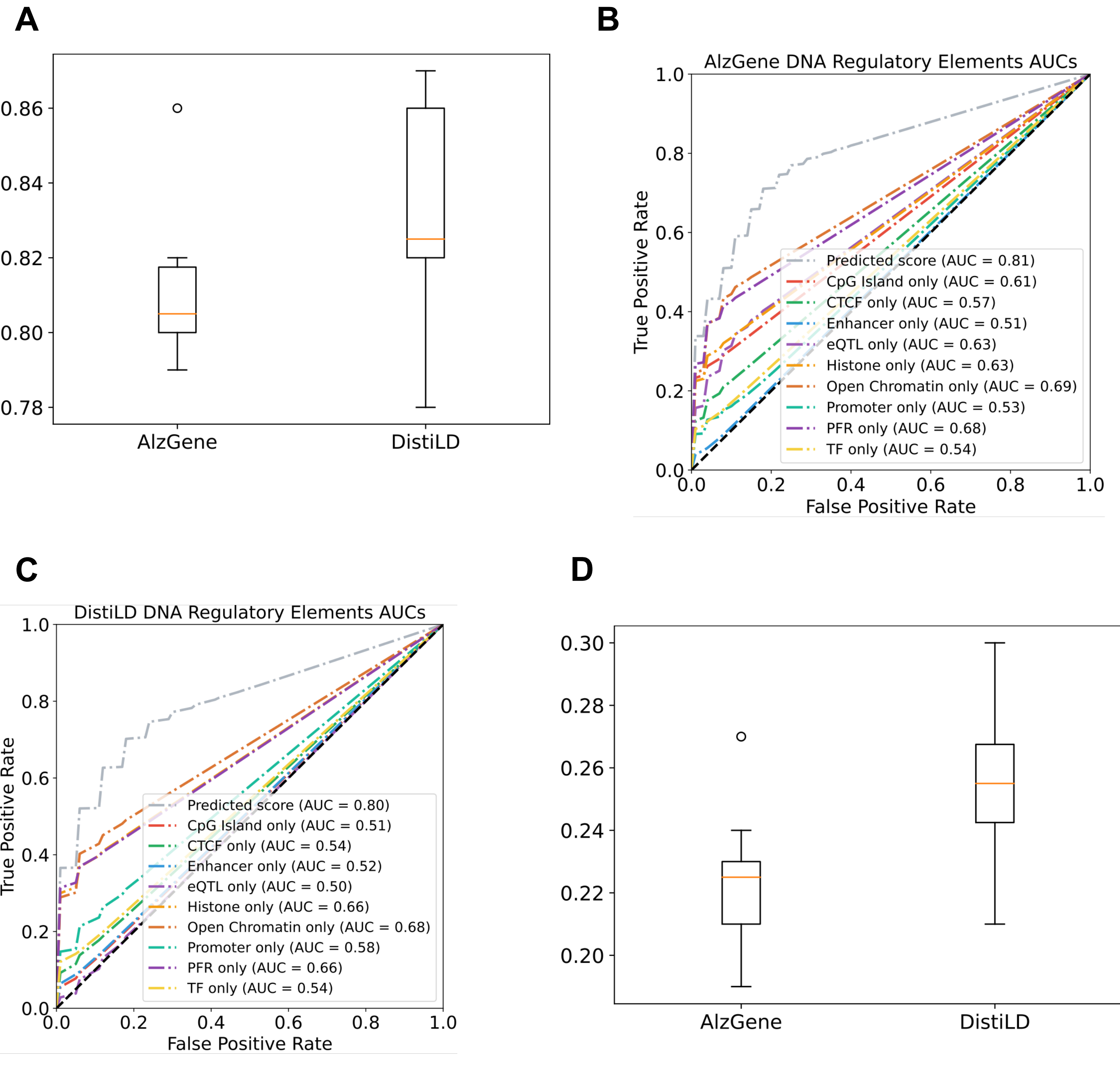

### Supplemental Figure S3

## A

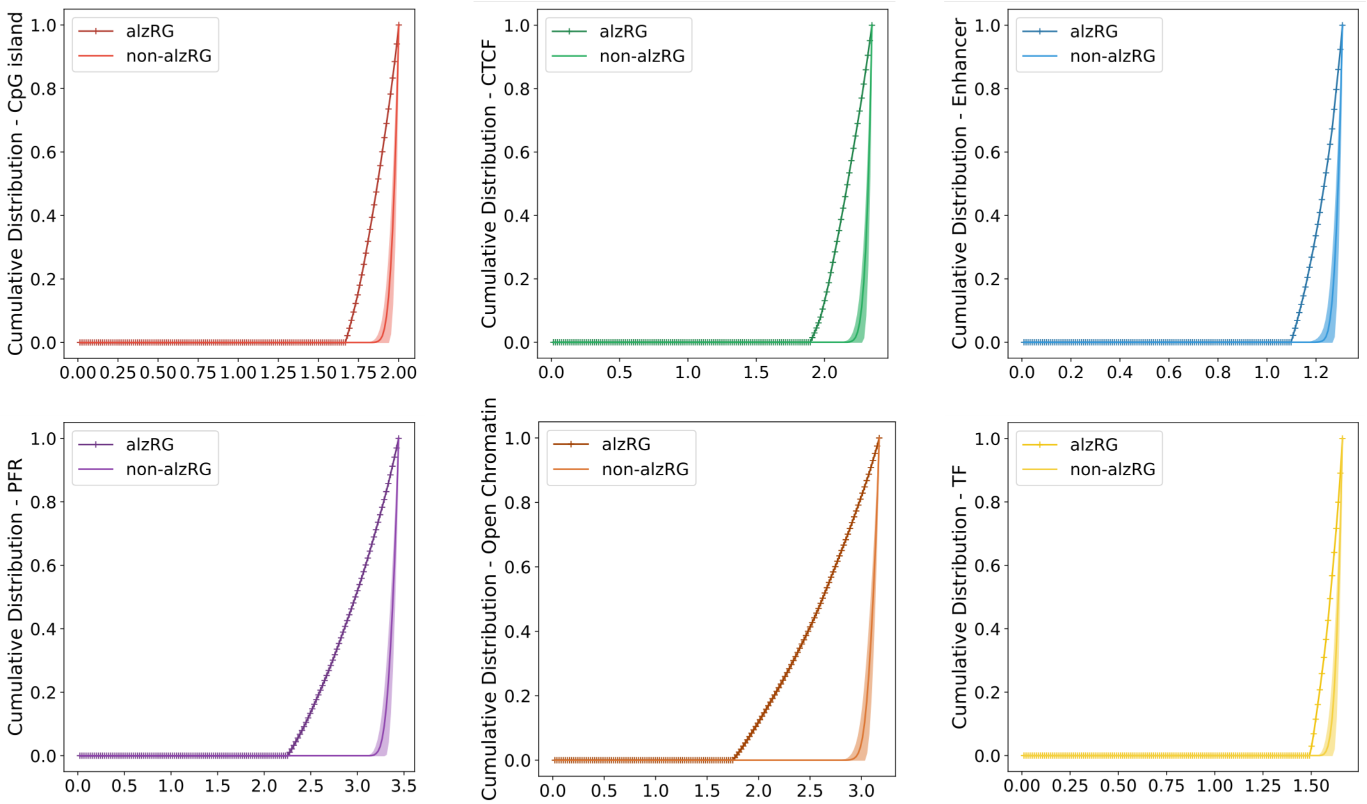


## B

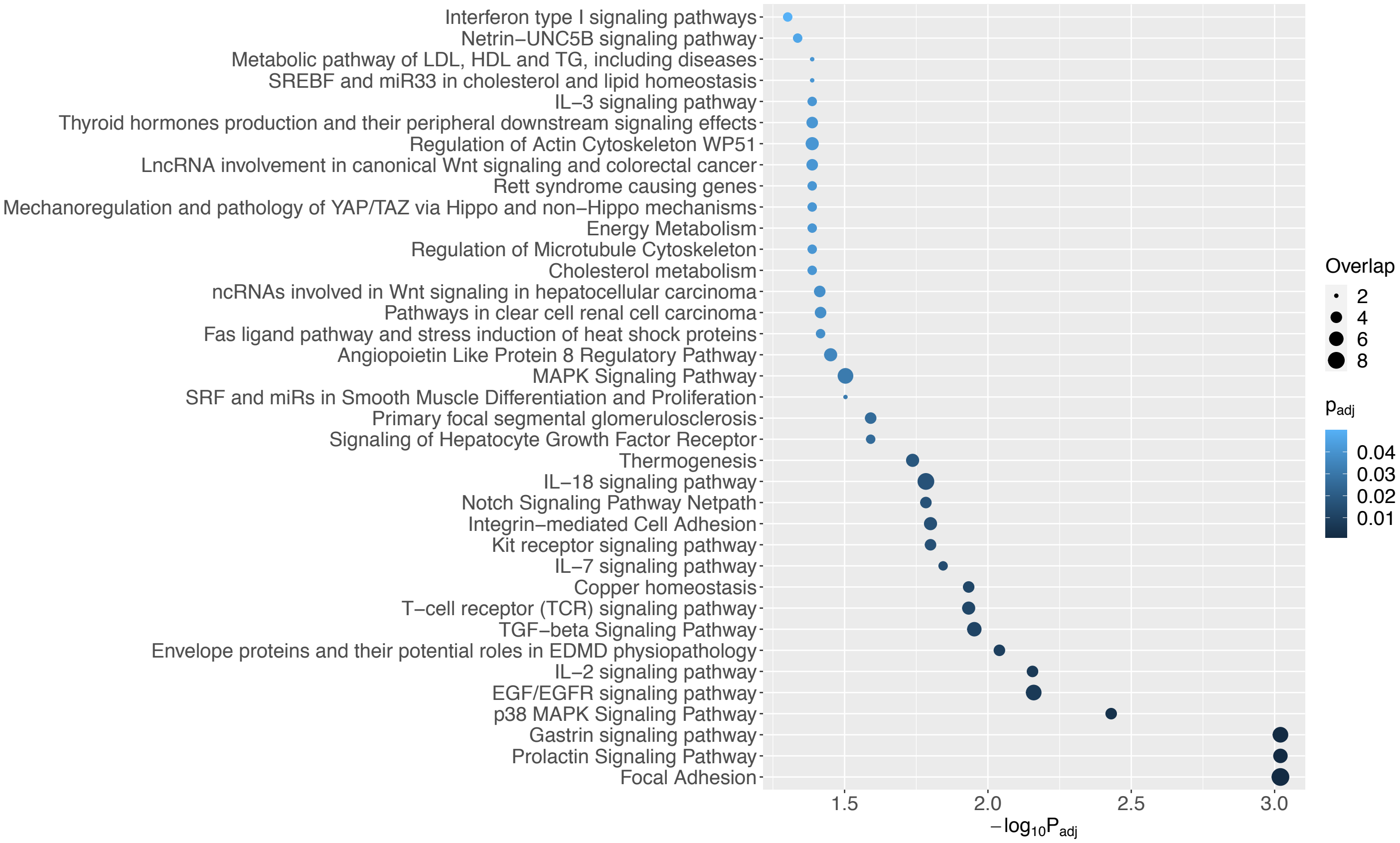

### Supplemental Figure S4

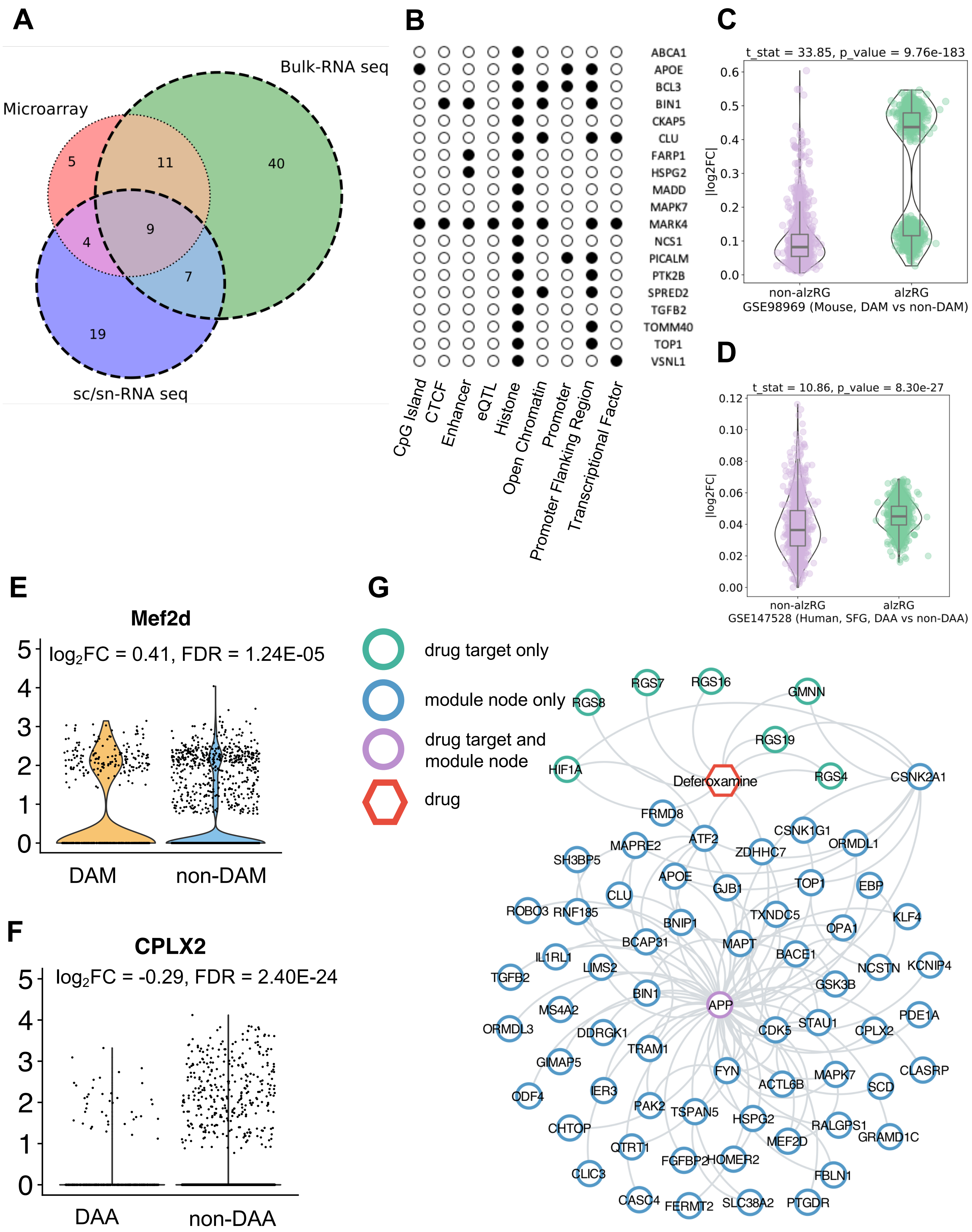

### Supplemental Figure S5

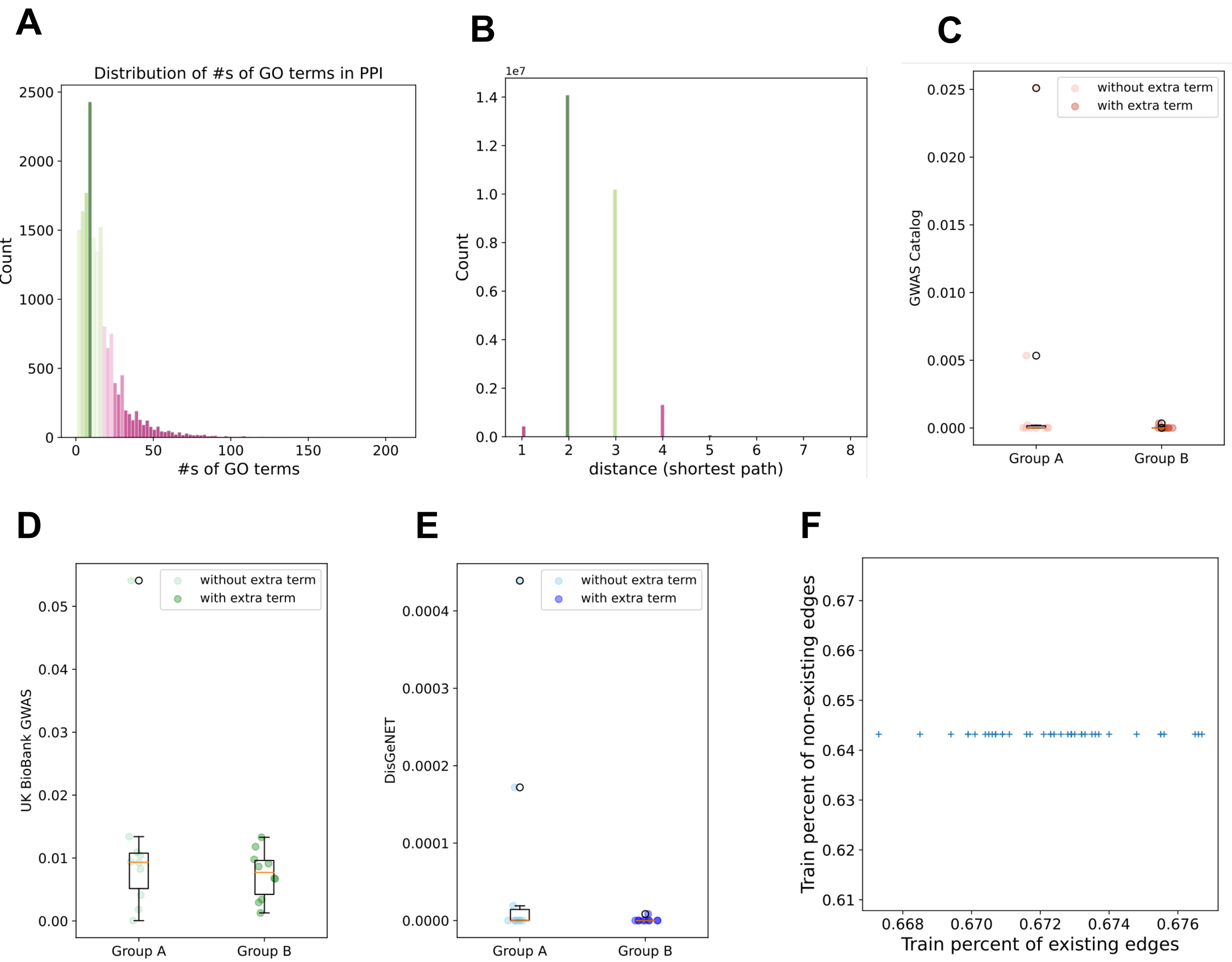
